# Immunokinetic Model for COVID-19 Patients

**DOI:** 10.1101/2022.01.13.476252

**Authors:** Y. Fadaei, F. A. Rihan, C Rajivganthi

**Author notes:** Corresponding should be addressed to (Y. Fadaei) & (F.Rihan).

## Abstract

In this paper, we develop a fractional-order differential model for the dynamics of immune responses to SARS-CoV-2 viral load in one host. In the model, a fractional-order derivative is incorporated to represent the effects of temporal long-run memory on immune cells and tissues for any age group of patients. The population of cytotoxic T-cells (CD8^+^), natural killer (NK) cells and infected viruses are unknown in this model. Some interesting sufficient conditions that ensure the asymptotic stability of the steady states are obtained.

This model indicates some complex phenomena in COVID-19 such as “immune exhaustion” and “Long COVID”. Sensitivity analysis is also investigated for model parameters to determine the parameters that are effective in determining of the long COVID duration, disease control and future treatment as well as vaccine design. The model is verified with clinical and experimental data of 5 patients with COVID-19.

## 1 Introduction

The ongoing pandemic coronavirus (CoV) disease outbreak (COVID-19) started in Wuhan, China, in December 2019, and has spread to more than 197 countries. Rapid spread of this disease threatens the health of a large number of people. As a result, immediate measures must be taken to prevent the disease in the community. It is the seventh member of the Coronavirus (CoV) family, along with MERS-CoV and SARS-CoV [1]. The virus is very serious and spread through respiratory droplets and close contact [2]. Scientists and researchers are therefore interested in how to develop treatment methods for such infectious diseases. Those methods are useful in understanding the dynamics/interactions between pathogens and their hosts. For years, mathematical modelers have been addressing specific aspects of infectious diseases [3, 4]. The majority of these efforts have been focused on multi-level diseases and have adopted quite different computational approaches [5, 6, 7, 8, 9].

Humans may develop upper-respiratory-tract infections as a result of COVID-19 transmission at the cellular level. Human cells have healthy, infected, virus cells and antibodies that are input parameters, and the output will be infected lung cells. The transmission of CoV among groups has been discussed in many research papers [10, 11, 12]. Despite this, the dynamics of CoV infection in an individual (organism) [22] are not extensively explored in the literature, which we analyze in the present paper.

In epidemiology and immunology, mathematical models are used to understand the dynamics of infectious diseases. In general, the coronavirus model depends on the initial conditions, and the classical order model cannot explain the virus perfectly because of its local nature.

Fractional-order derivatives are non-local in nature and are also dependent on the initial values. Furthermore, the fractional-order model has more advantages in terms of best fitting data, information about its memory, and hereditary properties. Furthermore, the hereditary properties increase the utility of the models constructed in fractional-order derivatives to describe the real phenomenon (see [13, 14]). In [15] studied the transmission dynamics of fractional-order coronavirus models and compared our results with some real data against confirmed infection and death cases per day for the first 67 days in Wuhan. According to [16], the authors compared the results of integer and fractional-order coronavirus (SEIRD) models, using real data from Italy, reported by the WHO. The results proved that the fractional-order case has a less root-mean-square error of fitting the model to the real data than the classical one and the fractional model has a closer estimation of the reality. Singh *et al.* [17] discussed the discretization computational techniques to solve numerically a fractional-order coronavirus model and this technique are effective to show the behavior of the solution in a very long time period which is helpful to predict the coronavirus model accurately. Most of the authors studied the coronavirus dynamics in the sense of fractional-order derivatives ([18, 19, 20, 21]). In cells level, the authors in [22] studied the dynamics of a fractional-order delay differential model for coronavirus (CoV) infection to give us best understand what causes the intensity of symptoms and illness of contaminated lung and respiratory system; See also [23, 24, 25].

As a result of the above motivation, in this paper, we propose a fractional-order model for coronaviruses with three compartments, such as SARS-CoV-2 density, cytotoxic T-cells, and natural killer cells. The Caputo fractional derivative has a power-law kernel, where its decaying rate depends directly on the fractional-orders. For the considered model, we derive the positiveness of the solution and examine the local stability of existing equilibrium points. By using the important sensitive parameters, we study the model qualitatively to demonstrate the eradication of the disease. As graphs, we can show more interesting results and their theoretical and numerical justifications.

This paper is organized as follows: In Section 2, we propose a virus infection model and study the positivity solution and local stability results. In Section 3, we discuss parameter estimation. Section 4 provides numerical simulations to validate the obtained theoretical results. Section 5 provides sensitivity analyses. The conclusion is in Section 6.

## 2 The Mathematical Model

Herein, we develop a fractional order mathematical model for the immune system response to SARS-CoV-2 virus in COVID-19 patients. We consider the RNA SARS-CoV-2 viral load (S), a cell population of the innate immune system: natural killer (NK) cells, and a cell population of the adaptive immune system: cytotoxic (CD8^+^) T-cells (T). Also, we assume that *t* represents the variable time (day). The assumptions of the model are:

- The populatin of infected cells and the SARS-CoV-2 virus concetration are assumed same.
- The SARS-CoV-2 virus in the absence of an immune response grows logistically. That is based on fitting of the data in [26].
- The infected virus can be cleared by both NK and CD8^+^ cells [26, 27].
- The virus promotes an initial activation of NK and CD8^+^ cells in the beginning of the disease [28, 30].
- The total number of NK cells was decreased in patients after some number of encounters with with SARS-CoV-2 infection [30].

Based on above assumptions, the system of fractional differential equations for representing interactions of SARS-CoV-2 virus and immune system is given by:

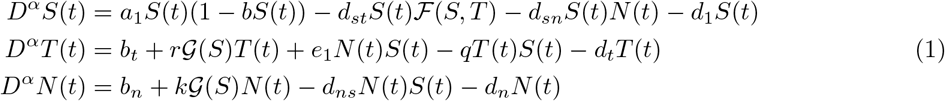

where 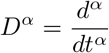, defined in Caputo sense. In the equations, three cell population are denoted by:

S(t) = density of SARS-CoV-2 (copies/*ml*),
T(t) = total cytotoxic T-cells population (cell/*ml*),
N(t) = total natural killer cells (cell/*ml*).

The term 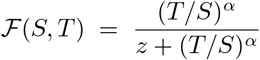 is fractional viral clearance rate of rational form by activated cytotoxic T-cells which is based on de Pillis–Radunskaya Law [36]. However, 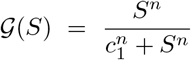 is a modified Michaelis-Menten term for T-cells activation and NK cell recruitment by SARS-CoV-2. *S*(0) = *S*_0_ > 0, *T*(0) = *T*_0_ > 0, *N*(0) = *N*_0_ > 0 are initial conditions of the system (1) and 0 < *α* ≤ 1 is derivative order.

The dynamics of the SARS-CoV-2 is represented by the first equation of system (1). Infected virus growth is logistically with replication rate *a*_1_ and carrying capacity *b*. Virus lysis by CD8^+^T-cells is shown by 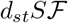, the term *d_sn_SN* represents the virus death by NK cells. Viral clearance rate is presented by *d*_1_.

The second equation shows the dynamic of the CD8^+^ T-cells against infected virus. Birth and death of CD8^+^ T-cells are represent by *b_t_* and *d_t_T* terms [31]. The term 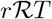 shows amount of CD8^+^ T-cells activation by infected virus. The term *e*_1_*NS* represents recruitment of CD8^+^T-cells by the debris from virus lysed by NK cells [32]. Inactivation of CD8^+^ T-cells by infected virus is shown by *qTS* term. Behaviar of NK cells are represented by third equation. NK cells activation by SARS-CoV-2 is shown by 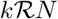. The term *d_ns_NS* is inactivation terms of NK cells by infected cells. Natural death of NK cells is represented by *d_n_N* term.

### Definition 2.1.

*[14] Caputo derivative of fractional-order α for a function f (t) is described as*

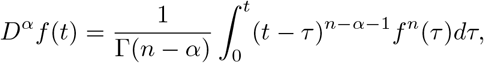

*where n – 1 < α < n ∈ ℤ ‘, Γ(·) is the Gamma function.*

The Laplace transform of Caputo derivative is described as:

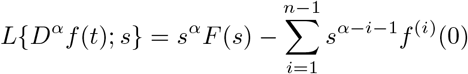

where *F*(*s*) = *L*{*f*(*t*)}. In particular, when *f*^(*i*)^(0) = 0, *i* = 1, 2,..., *n* – 1, then *L*{*D^α^f* (*t*); *s*} = *s^α^F*(*s*).

The basic reproductive rate/ratio, 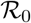 is defined as the expected number of secondary infections arising from a single individual during his or her entire infectious period, in a population of susceptibles. Epidemiology and pathogen dynamics within hosts are both based on this concept. Furthermore, 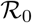 is used as a threshold parameter that predicts whether an infection will spread. However, related parameters that share this threshold behavior may or may not give the true value of 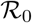. It also denotes as the number of secondary infection due to a single infection in a completely susceptible population. We derive the expression of 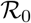, allied to the disease-free equilibrium 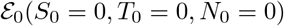. The recovery rate from the virus, and transmission rate of the virus from infected individuals to susceptible individuals are described by the following matrices

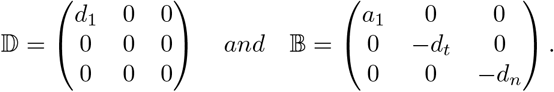

Thus, the basic reproduction number 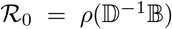, calculated as the spectral radius of the next generation matrix [33], is then defined by

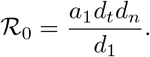

The disease is eradicated if 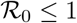 and will persist as *t* goes to infinity if 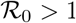; See [34].

### 2.1 Non-negativity of the model solutions

Herein, we investigate the non-negativity of the model solutions.

#### Lemma 2.2.

*(Generalized Mean Value Theorem [35]) Let the function 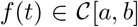 and its fractional derivative 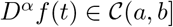 for 0 < a ≤ 1 and a,b ∈ ℝ then we have*

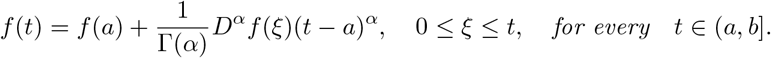

#### Remark 2.3.

*Assume that the function f (t) is α-differentiable on (a, b) then we have the following results [14]:*

- *If D^α^ f (t) < 0 for all t ∈ (a, b) then f (t) is decreasing on (a, b).*
- *If D^α^ f (t) > 0 for all t ∈ (a, b) then f (t) is increasing on (a, b).*
- If D^α^ f (t) = 0 *for all t ∈ (a, b) then f (t) is constant on (a, b).*

#### Lemma 2.4.

*The solutions of model (1) with nonnegative initial values are non-negative.*

*Proof.* To show this Lemma, we ought to consider that the domain Ω = {(*S, T, N*) ∈ ℝ^3^: *S* ≥ 0, *T* ≥ 0, *N* ≥ 0} is positively invariant region. Then, on the hyperplanes of region Ω we have

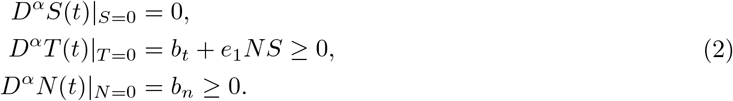

If {*S*(0), *T*(0), *N*(0)} ∈ Ω according to Lemma 2.2 and Remark 2.3, the solution (S(t), T(t), N(t)) can not escape from the hyperplanes Ω. Thus, the solutions of the fractional-order model (1) are non-negative if the initial conditions are non-negative for all *t* > 0. ?

### 2.2 Stability of the steady states

The underlying model (1) has the following equilibrium points: i) Decease free with immunity equilibrium 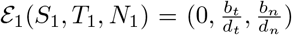, ii) Endemic equilibrium point 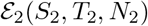, if they exist, satisfy the following equalities:

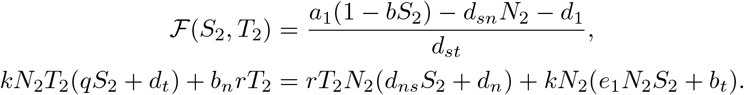

The corresponding linearized system of model (1) at any steady state (*S*\*,*T*\*,*N*\*) is calculated as follows

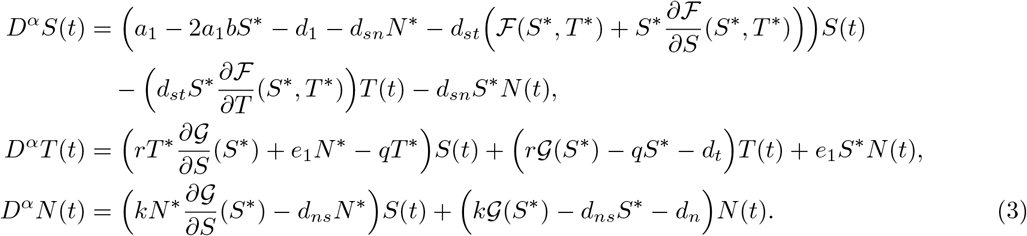

Applying Laplace transform on both sides of (3), we can get

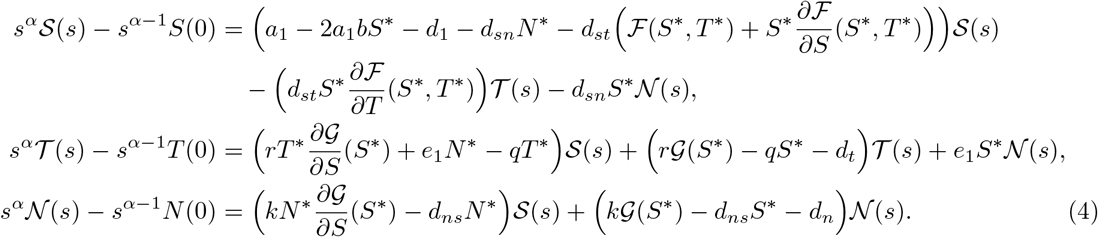

Here, 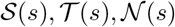 are Laplace transform of *S*(*t*), *T*(*t*) and *N*(*t*) with 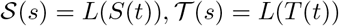 and 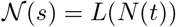. The above equations (4) can be written as

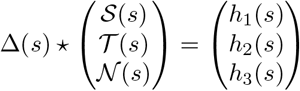

where

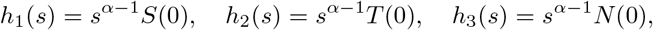

and

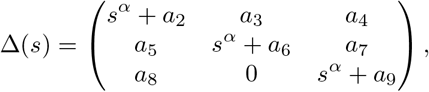

Δ(*s*) is the characteristic matrix for the system (3) at (*S*\*,*T*\*,*N*\*) and

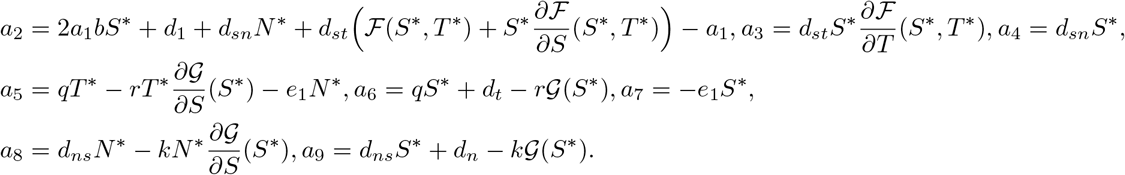

Clearly, the eigenvalues of Δ(*s*) at 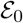 and 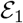 are -*d_t_*, -*d_n_*, *d*_1_ — *a*_1_ and -*d_t_*, -*d_n_*, *d*_1_ + *d_sn_N*_1_ – *a*_1_, respectively, and assume that *d*_1_ < *a*_1_,*d*_1_ + *d_sn_N*_1_ < *a*_1_, which confirm that the model (1) around the equilibrium points 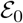 and 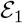 are stable.

#### Lemma 2.5.

*The endemic equilibrium point 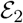 is locally asymptotically stable if p_1_ > 0, p_3_ > 0, p_1_ p_2_ > p_3_.*

*Proof.* The characteristic equation at 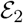 is described by

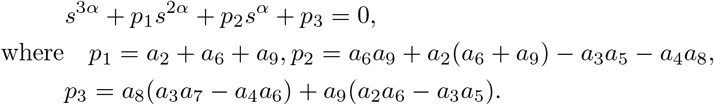

By using the Routh-Hurwitz criterion, the endemic equilibrium 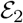 is locally asymptotically stable if *p*_1_ > 0, *p*_3_ > 0, *p*_1_*p*_2_ > *p*_3_.

## 3 Parameter Estimation

The study of Wölfel *et al.* [26] was done a virological analysis on nine patients of COVID-19 for examining the kinetics of viral load and measuring the virus replication in tissues of the upper respiratory tract. Infection of all patients was known because they had near contact to an index case. The patients were admitted to a hospital in Munich, Germany, and underwent virological tests in collaboration with two reputable laboratories. Both laboratories were equipped with the same technology in PCR-PT and the same standards for virus isolation. Authors measured and analyzed viral loads were projected to RNA copies per *ml*, per *swab* and per *g* for sputum, throat swab and stool samples, respectively. All samples were taken between 2 and 4 days after the onset of symptoms. In [29] swab samples are used for some mathematical models.

Here, data fitting is used to estimate the values of parameters of the model (1). The parameters are fitted by measured RNA viral load in sputum samples of five patients from [26] using by implementing a least squares algorithm, fminsearch, that is a MATLAB function. The measured viral load was done daily. The results for parameter estimation are presented in Table 1. Data fitting are made for different valuse of *α* ∈ (0,1]. In Figures 1–5 the result of the fitting for values *α* =1 and *α* = 0o98 are presented. Due to the arbitrary derivative order of model and non–locality properties of these derivatives, different curves may be obtained in data fitting. This advantage will help to find the best fitting to the parameters of the model.

**Figure 1:**
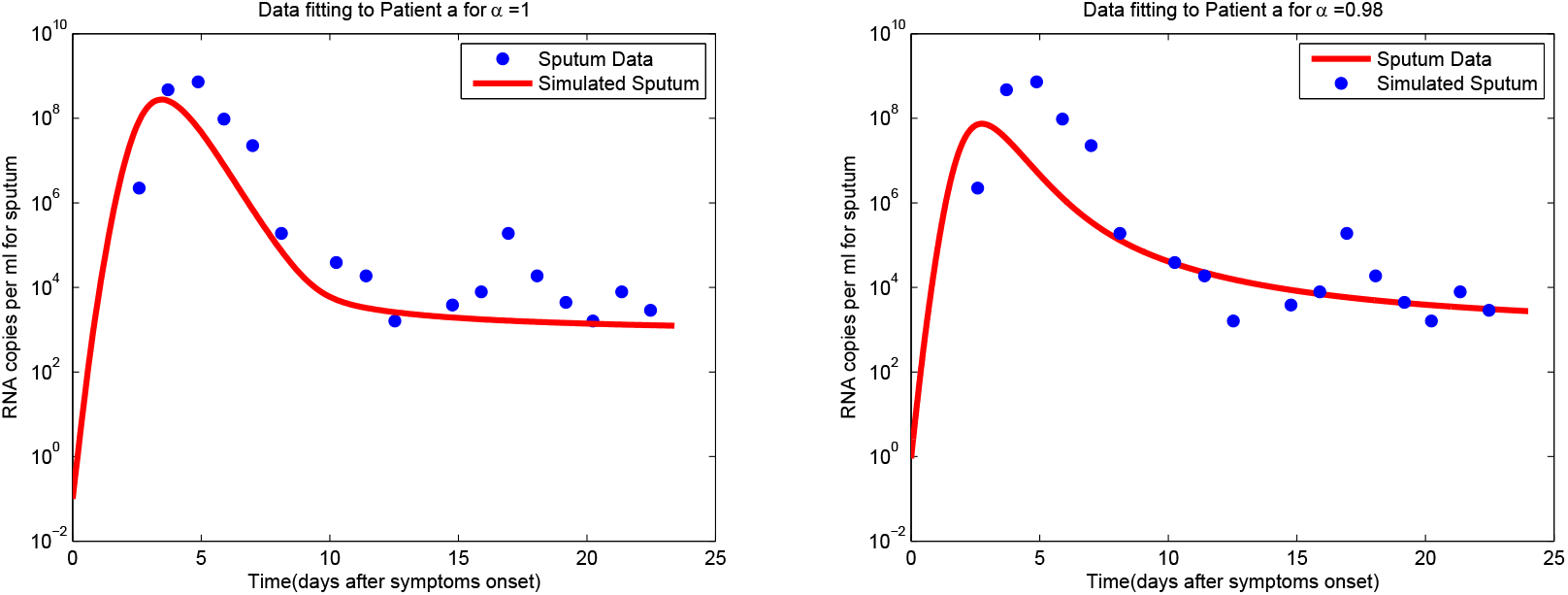
Data fitting for patient **a**.

**Figure 2:**
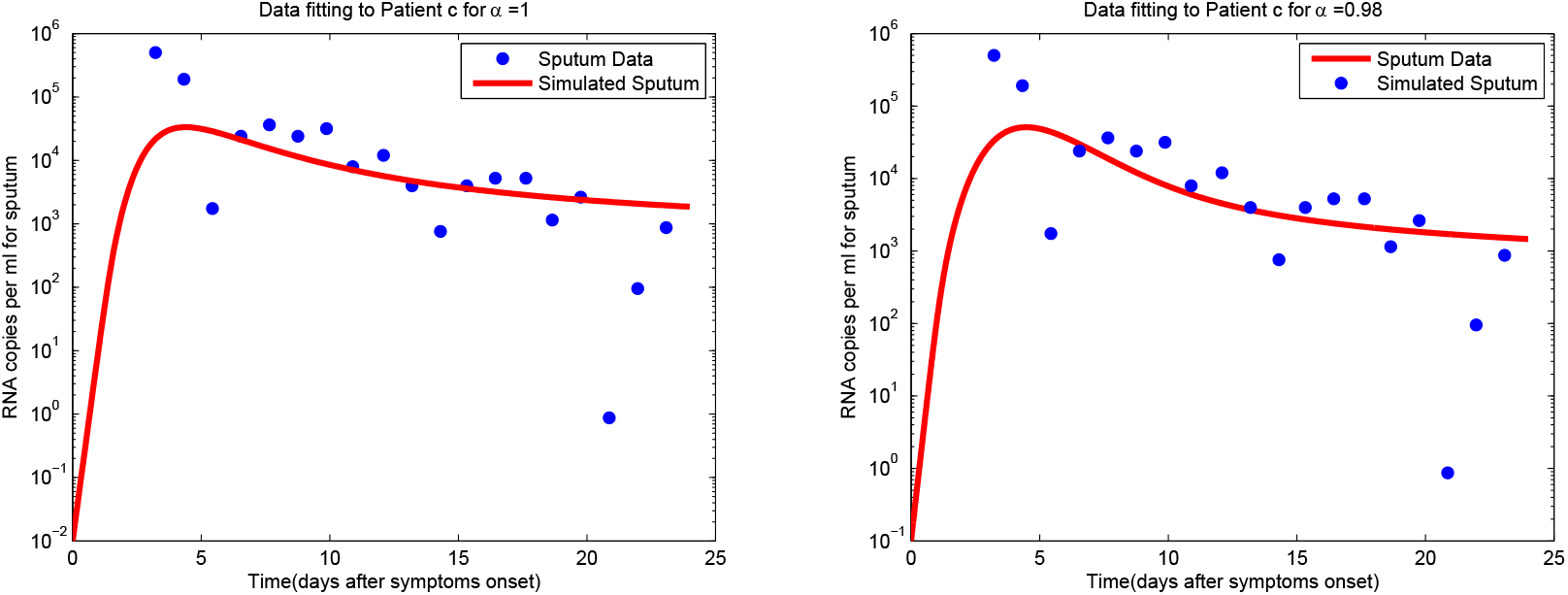
Data fitting for patient **c**.

**Figure 3:**
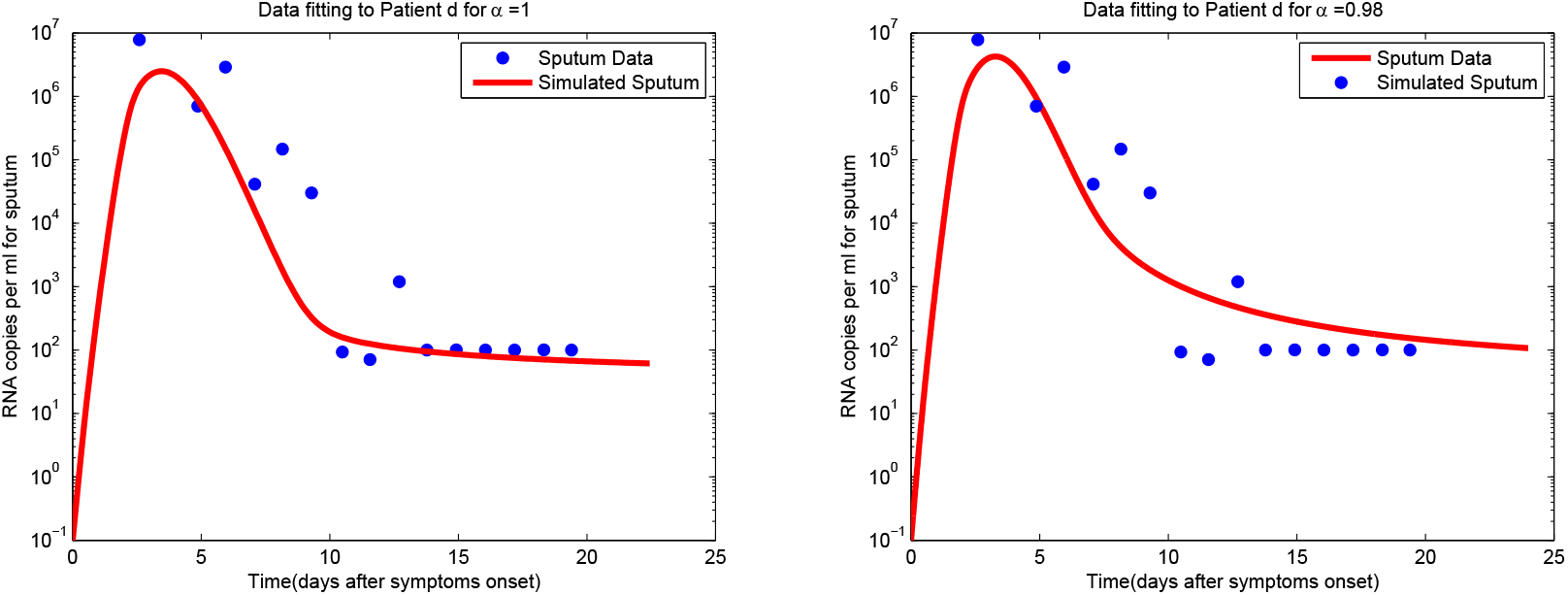
Data fitting for patient **d**.

**Figure 4:**
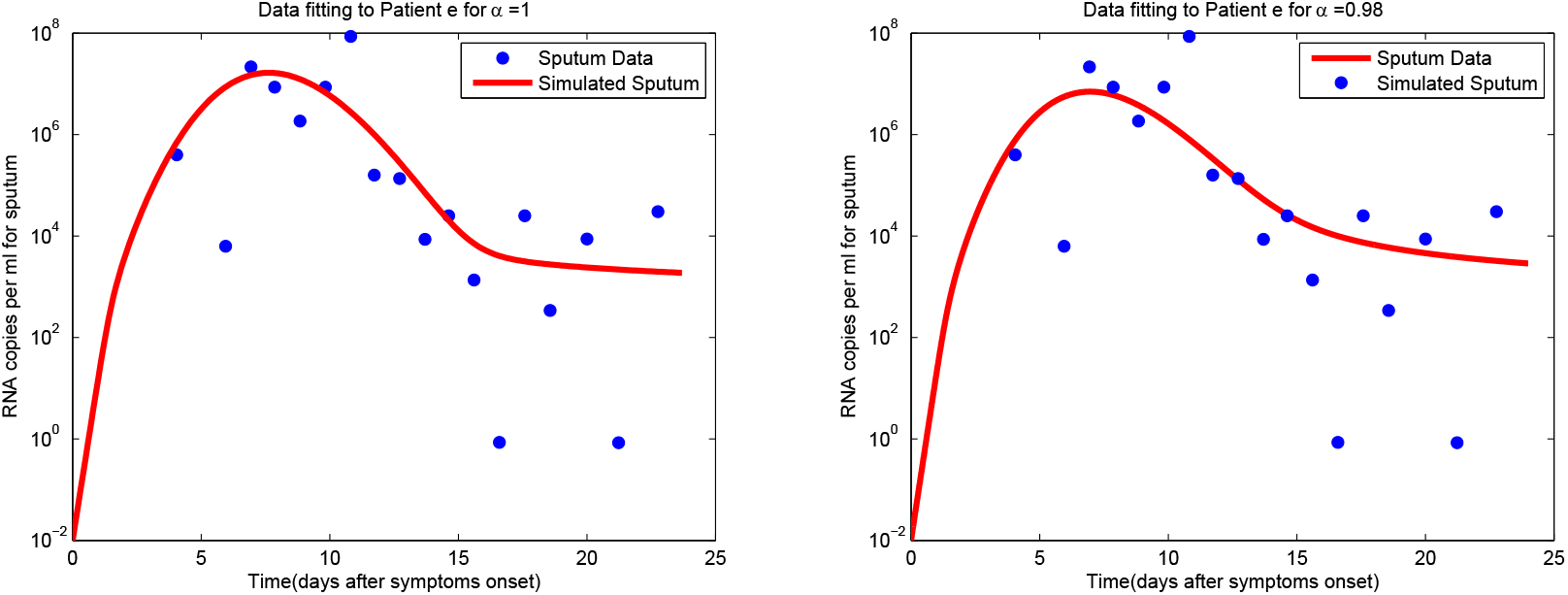
Data fitting for patient **e**.

**Figure 5:**
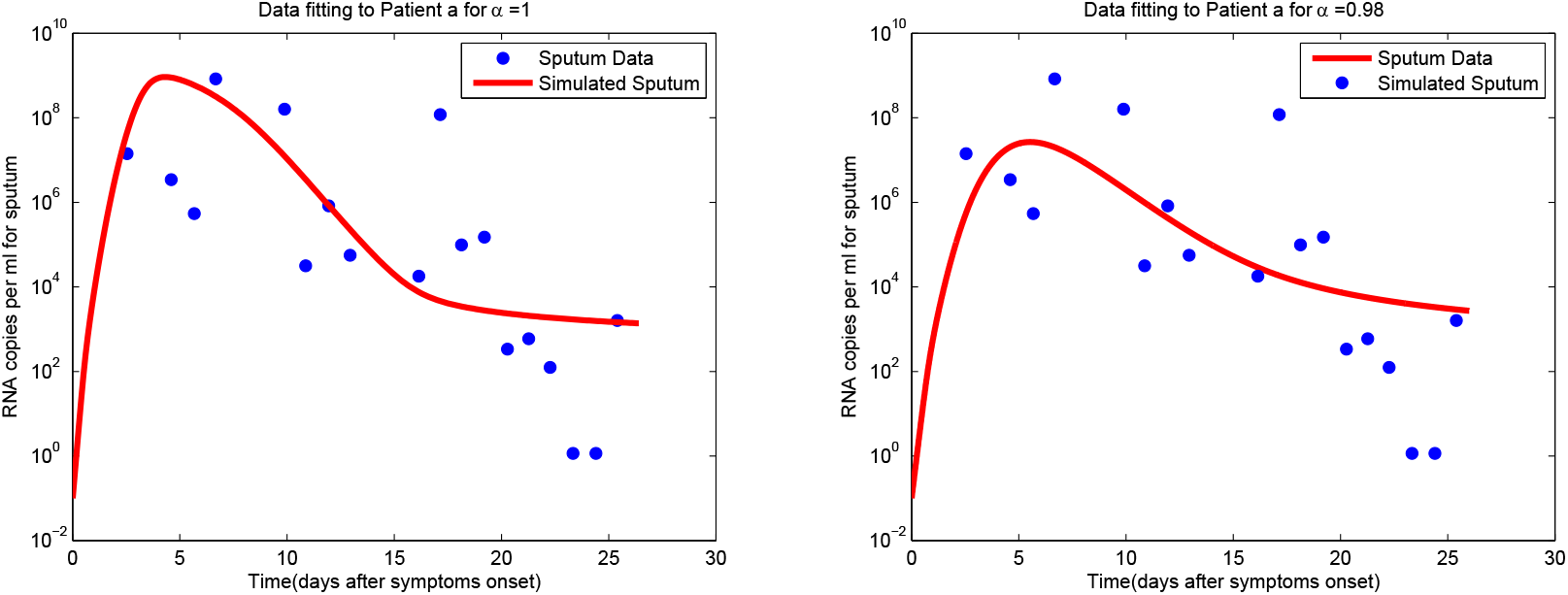
Data fitting for patient **g**.

**Table 1:** Information of the Parameters.

| Par. | Description | Units | Value range | Source |
| --- | --- | --- | --- | --- |
| $a_1$ | S replication rate | 1/day | [2.86,7.07] | [29] |
| $b$ | Maximum S | ml/(RNA copies) | [1e-010,1e-09] | [29] |
| $d_{st}$ | Virus lysis by CD8<sup>+</sup> T-cells rate | 1/day | [0.01,0.4] | Estimated |
| $d_{sn}$ | Virus death rate by NK cells | ml/cell(day) | [1.2e-11,2e-010] | Estimated |
| $d_1$ | Natural death rate of Virus | 1/day | [0.0001,0.1] | Estimated |
| $b_t$ | CD8<sup>+</sup> T-cells proliferation | cell/ml(day) | [50 1500] | [47] |
| $r$ | CD8<sup>+</sup> T-cell activation | 1/day | [0.001,0.032] | Estimated |
| $e_1$ | CD8<sup>+</sup> T recruitment rate by virus lysed by NK | ml/(RNA copies)(day) | [2.1e-06,8.4e-005] | Estimated |
| $q$ | CD8<sup>+</sup> T inactivation rate by virus | ml/(RNA copies)(day) | [1.1e-10,9.5e-10] | Estimated |
| $d_t$ | Natural death rate of T-cells | 1/day | [0.001,0.08] | [48] |
| $b_n$ | NK cells proliferation | cell/ml(day) | [7, 200] | [43] |
| $k$ | NK cells activation | 1/day | [0.001,0.02] | Estimated |
| $d_{ns}$ | Inactivation NK cells rate | ml/(RNA copy)(day) | [1.1e-06,9.09e-05] | Estimated |
| $d_n$ | NK cells death rate | 1/day | [4.2e-02,0.15] | [46, 45] |
| $z$ | Steepness coefficient of virus lysis by T-cells | — | [0.01,1] | [43] |
| $c_1$ | Steepness coefficient of NK recruitment | — | [1e+03,1e+06] | Estimated |
| $\alpha$ | Fractional virus kill power | — | [9.1e-01,9.9e-01] | Estimated |
| $n$ | Michaelis-Menten order | — | 2 | [48] |

## 4 Simulations and Model Validation

In order to numerically solve the system (1) the Adams-Bashforth-Moulton method of fractional version (FABM) will be used. This method was introduced in [37]. Consider following fractional-order differential equation

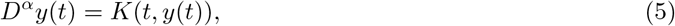

the fractional Adams-Bashforth-Moulton method is include two step first step is predictor:

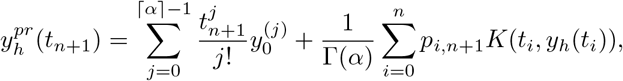

after computing the predictor step in second step modifier is calculated by

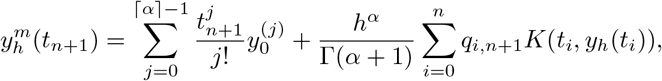

where the *p*_*i,n*+1_ and *q*_*i,n*+1_ are

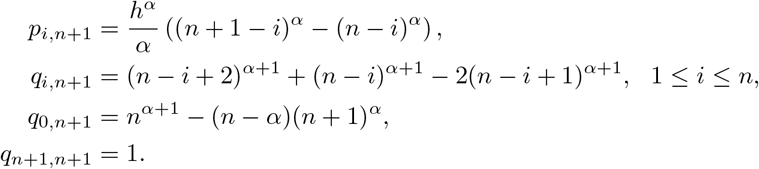

in which *t_i_ i* = 0,1...*n* are equally selected points with fixed step length *h*.

Garrappa has written a MATLAB function for FABM, FDE12, which is available at the MathWorks [50]. The FDE12 algorithms is used for numerically solving of the model (1). This numerical simulation is done for five patients **a, c, d, e, g** in [26] with their associated parameter values.

All simulations were performed to evaluate the behavior of the SARS-CoV-2 virus against immune cells 28 days after the onset of symptoms. In these simulations, three values *α* = 0.95,0.85, 0.80 are considered as the fractional-order derivatives order of the equations in the model (1). The results are shown in Figures 6–8.

**Figure 6:**
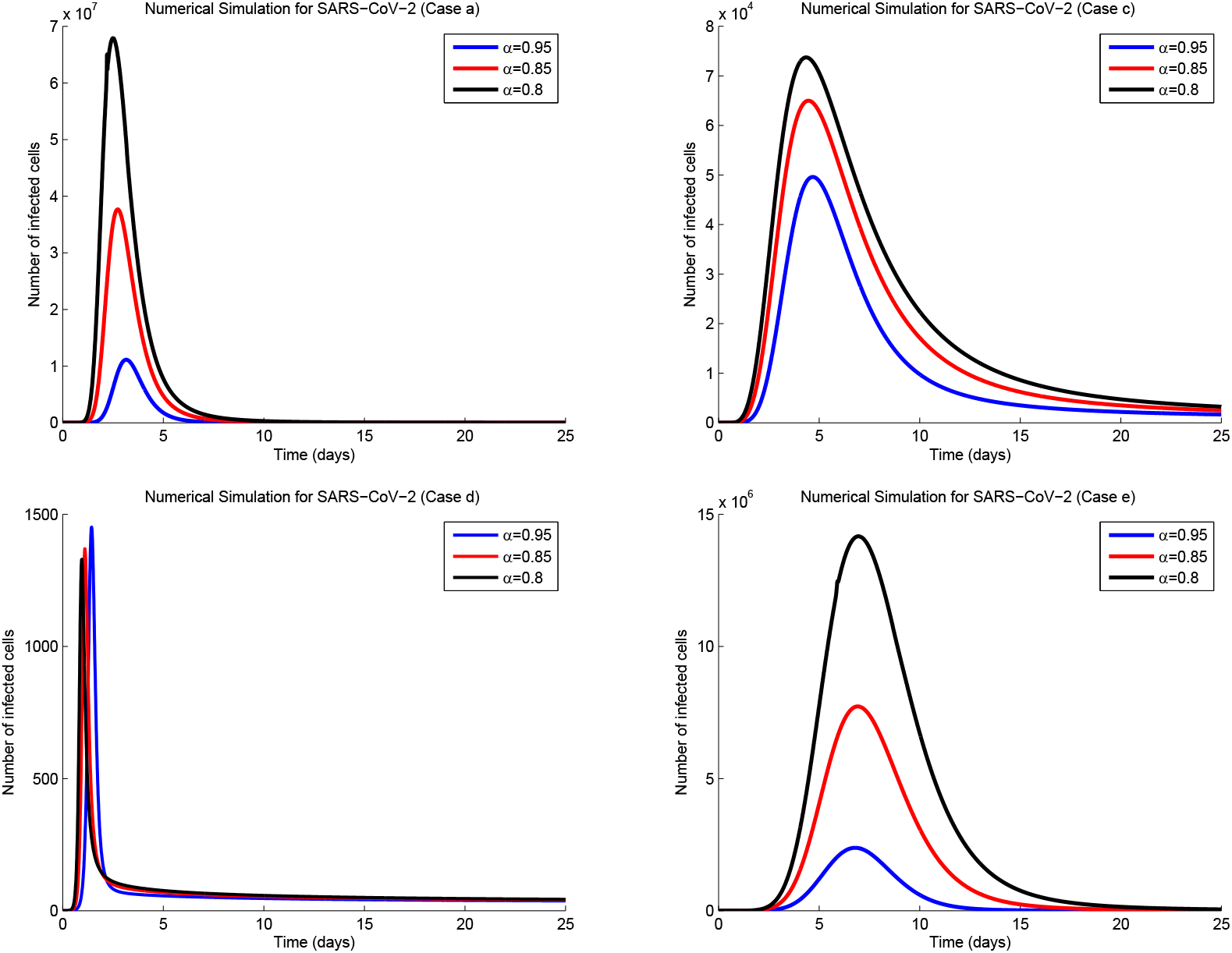
Numerical simulation results of SARS-CoV-2 behavior in patients **a**, **c**, **d**, **e**.

**Figure 7:**
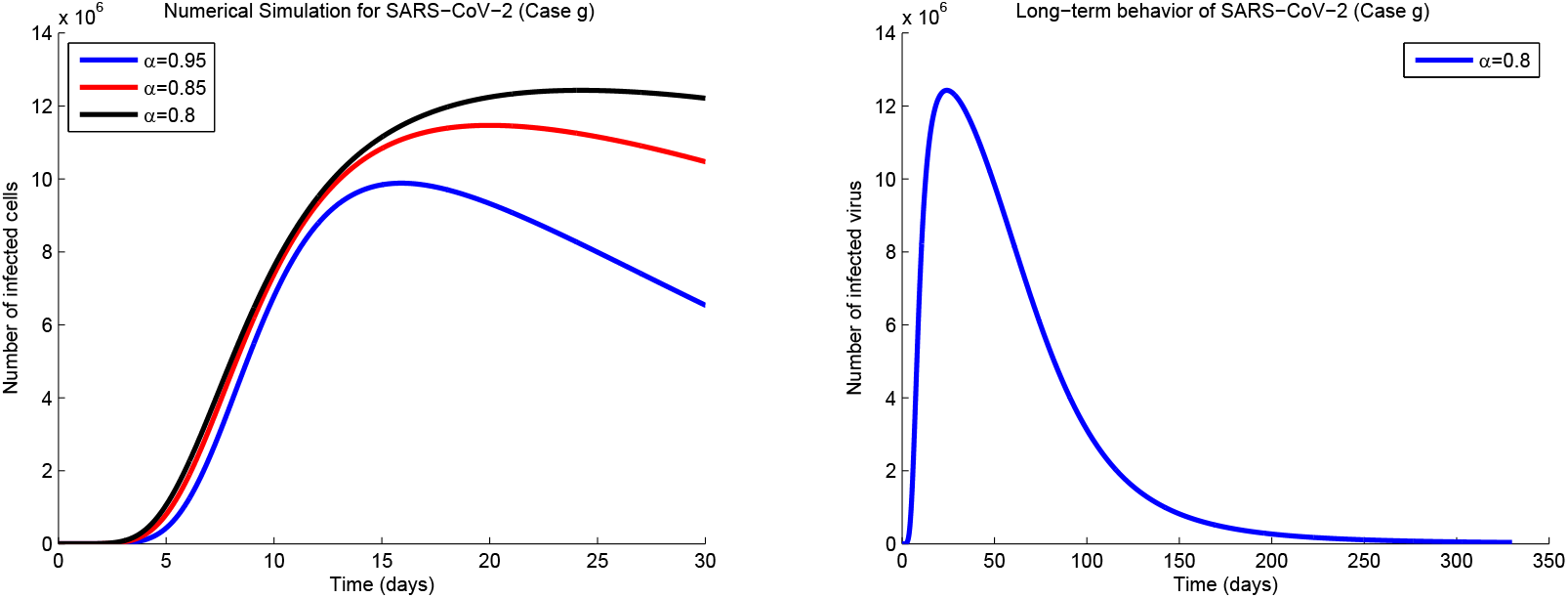
Numerical simulation results of SARS-CoV-2 behavior in patients **g** (left). Schematic of SARS-CoV-2 long-term behavior in patient **g** (right).

**Figure 8:**
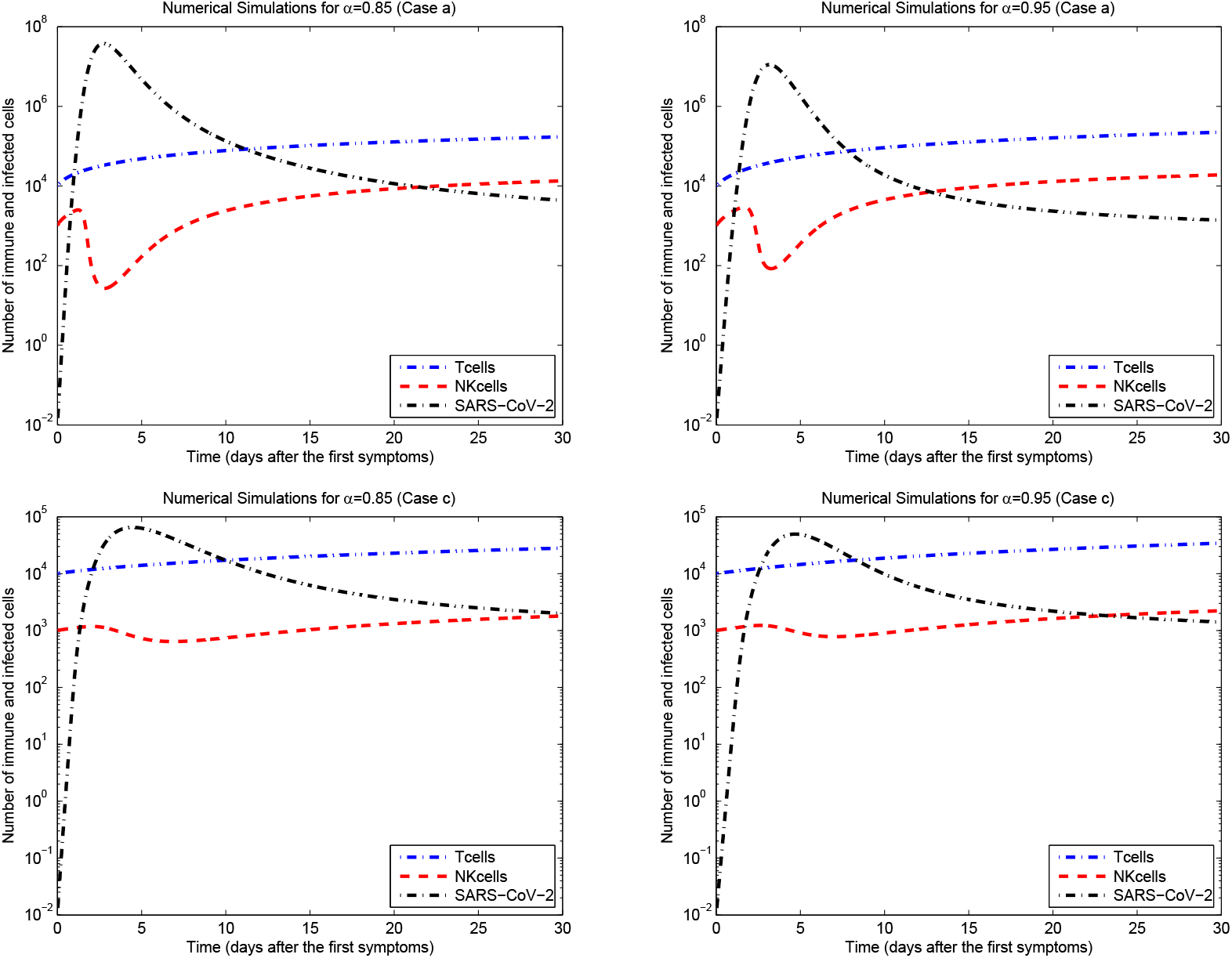
Numerical solutions of the model (1) for patient **a**, **c**.

As can be seen in the graphs, the virus concentration is not accurate for each patient. Simulations show that for smaller amounts of α the virus load is higher. It will also reduce the virus load with less speed and longer time.

In three patients **a, c** and **d**, the maximum load of the virus is before the fifth day. This may depend on the amount of contact and the amount of primary virus that has been transmitted to the patient. Of course, the initial behavior of the patient’s immune system against the virus should not be ignored.

Unlike patients **a, d**, in patient **c**, the decrease in the RNA viral concentration to its lowest level is about 30 days after the onset of symptoms. Slow in reducing the concentration of viral RNA due to the phenomenon of *immune exhaustion* and especially here *NK cell exhaustion*. High infections usually lead to NK cells exhaustion, so limiting the infection potential of NK cells [28, 27]. In SARS-CoV-2 infections exhaustion of the NK cell was confirmed by increased frequencies of programmed cell death protein 1 (PD-1) positive cells and reduced frequencies of natural killer group 2 member D (NKG2D)-, sialic acid-binding Ig-like lectin 7 (Siglec-7)-, and DNAX ancillary molecule-1 (DNAM-1)-expressing NK cells related to a reduced ability to spatter interferon IFN_γ_ (see Figure 9) [27]. Furthermore, it was shown that in sera of COVID-19 patients, IL-6 is present in large surplus. It may down-regulate NKG2D on NK cells, leading to disorder of NK cells activity [27].

**Figure 9:**
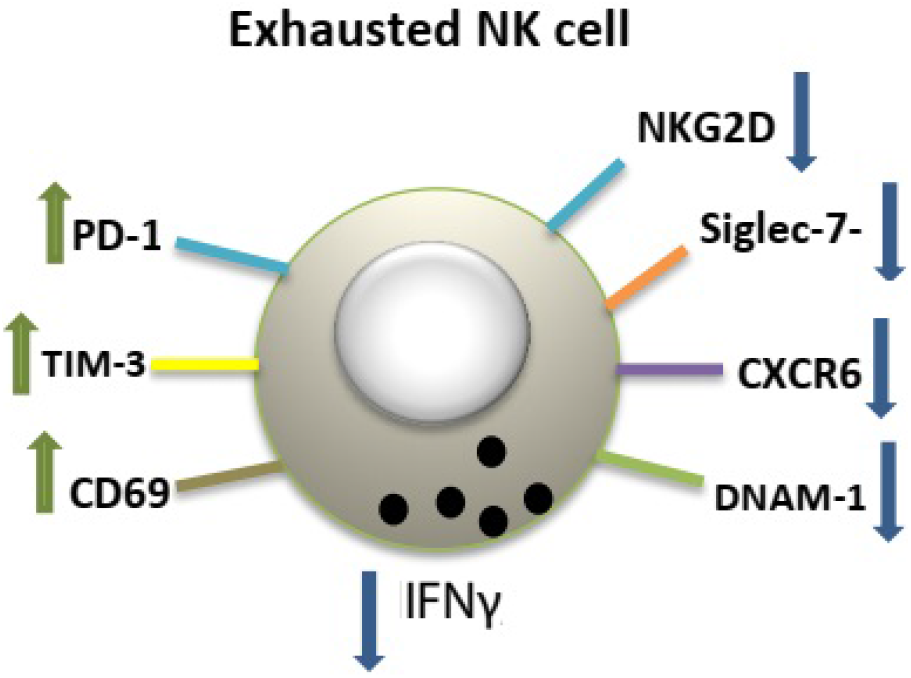
Schematic of exhausted NK cell.

In middle-aged patient **g**, due to the increased load of the virus, it leads to the NK cells exhaustion and reduces the infection potential. Decreased immune system function prolongs the course of the disease, so these patients need long-term treatment and a longer quarantine period than other patients. In addition, patients who show high viral loads 10 to 11 days after the first symptoms, due to immune exhaustion, will have symptoms of a lung infection [27]. If the limit of quantification of RNA viral load be 200 RNA copies per *ml,* the concentration of the virus in the patient’s body will reach this limit after 330 days for *α* = 0.8 (see Figure 7, right). In this case, it is said that the patient is involved in *Long COVID* or *Post COVID* phenomenon. Chronic COVID, known in English as long COVID, is a long-term symptoms of acute COVID disease. The disease, which is characterized by long-term complications, persists after a normal recovery period. The diagnosis of the duration or how long these conditions last is not yet fully understood [38]. Based on our model duration of long COVID for patient **g** is 330 days. Of note, it seems that delay in vaccination of immune exhaustion and long COVID individuals may be necessary. In the next section, we will discuss the process of the disease profile in the patient **g**.

In patient **e**, an increase in viral load occurs after the first week, potentially indicating an exacerbation of symptoms [26]. The immune system function of these patients need further investigation and more studies should be done in future studies.

The diagrams in Figure 8 show the behavior of infected virus versus the behavior of NK and CD8^+^ cells one month after the first symptoms in **a, c** patients. Order derivative values *α* = 0.95,0.85 are considered for both cases. System (1) solutions with *α* = 0.85 indicate that in patient **a** because of severe NK cells depletion as the first defense factor, SARS-CoV-2 virus growth reaches more than 10^7^. Two weeks after the peak viral load and with more CD8^+^ T-cells activation, the NK cells population increases and dominates the SARS-CoV-2 virus population. The solutions of Model (1) for *α* = 0.95 show that after approximately 10 days from the peak of infected virus concentration, the population of NK cells increases and overcomes viruses. Therefore, it can be said that it takes two to three weeks for the immune system to completely overcome the disease, in the patient **a**.

For the patient **c**, the solutions of (1) with *α* = 0.95,0.85 indicate that due to the greater resistance of NK cells to increased virus load and activated T-cells, the virus concentration is a maximum of 10^5^. Compared to the patient **a**, RNA viral had a lower burden, but due to NK cell exhaustion, NK cells were able to dominated SARS-CoV-2 infection with greater delay. About 25-30 days after the onset of symptoms, NK cells can return to their original value and completely dominate the infected virus.

The results of [42] show that despite the same initial viral load, innate immunity such as NK cells and INF_γ_, are stronger in younger patients and are more active than in adults in exposure to SARS-CoV-2 and quickly return to homeostasis. This may be seen in the solutions of model (1): As shown in Figure 8, when the *α* value is closer to one, the NK cells proliferate and become active faster. Also, the model with smaller *α* values is suitable for older patients. Here we can call *α* as the *age parameter*.

The response of CD8^+^ T-cells to the COVID virus is slow and at a constant rate. It seems that in order to reduce the peak load of the virus, T-cells need to respond more quickly to the virus attack. Therefore, it is recommended that in the first days of the disease, drugs that lead to faster activation of T-cells be prescribed. Rapid production of neutralizing antibodies is effective in treating the disease. In patients who made the neutralizing antibody before day 14, they eventually recovered, but in patients who started making the neutralizing antibody after 14 days, the antibodies lost their protective role [41].

## 5 Sensitivity Analysis

Sensitivity analysis is an important tool for assessing dynamic behavior of the underlying biological system. Herein, we evaluate sensitivity of state variables to small variations in model parameters to enable us to (i) display how robustness of the underlying infection model is to small changes in the parameter values, (ii) discover in which subinterval the model sensitive to a particular parameter to understand significant processes and immune system mechanisms. We evaluate the sensitivity functionals throughout studying the effect of changes in the parameters on the period to estimate severity of the diseases [22].

Some model parameters are very effective in determining the progression and decline of SARS-CoV-2 load. To determine the relationship between the parameters and model outcomes we use sensitivity analysis. Here we use Partially Ranked Correlation Coefficients (PRCC) to quantify the sensitivity and the relationships. The PRCC will be calculated for 1000 values of each parameter which is drawn by running the Latin Hypercube Sampling method (LSH). The LSH technique is a type of Monte Carlo sampling described in [39]. The LHS scheme allows the values of all input parameters to be changed simultaneously. This sampling method will be efficient if the outcome is a monotonic function of each of the input parameters. Here, we only use the parameters *a*_1_, *d_sn_, d_t_,b_t_, d_n_,b_n_,T*_0_ and *N*_0_ that are monotically associated to outcomes of the model in the sensitivity analysis.

Sensitivity analysis of the selected parameters was performed for 4 and 23 days post-onset of symptoms. The results for SARS-Cov-2 load are peresented in Figure 10. On day 4 after the first symptoms, the parameter *a*_1_, which is replication rate of the virus had a significant positive relationship with virus load. The PRCC value for the parameter *a_1_* at significance level of 0.001 was 0.62. The virus lysis by CD8^+^ T-cells rate parameter *d_st_* had a high negative correlation with viral load. The correlation coefficient for this parameter was 0.87. This negative correlation with viral loading indicates that increasing the SARS-CoV-2 lysis by CD8^+^ T cells may play an important role in controlling and reducing the virus load in the first days of the disease.

**Figure 10:**
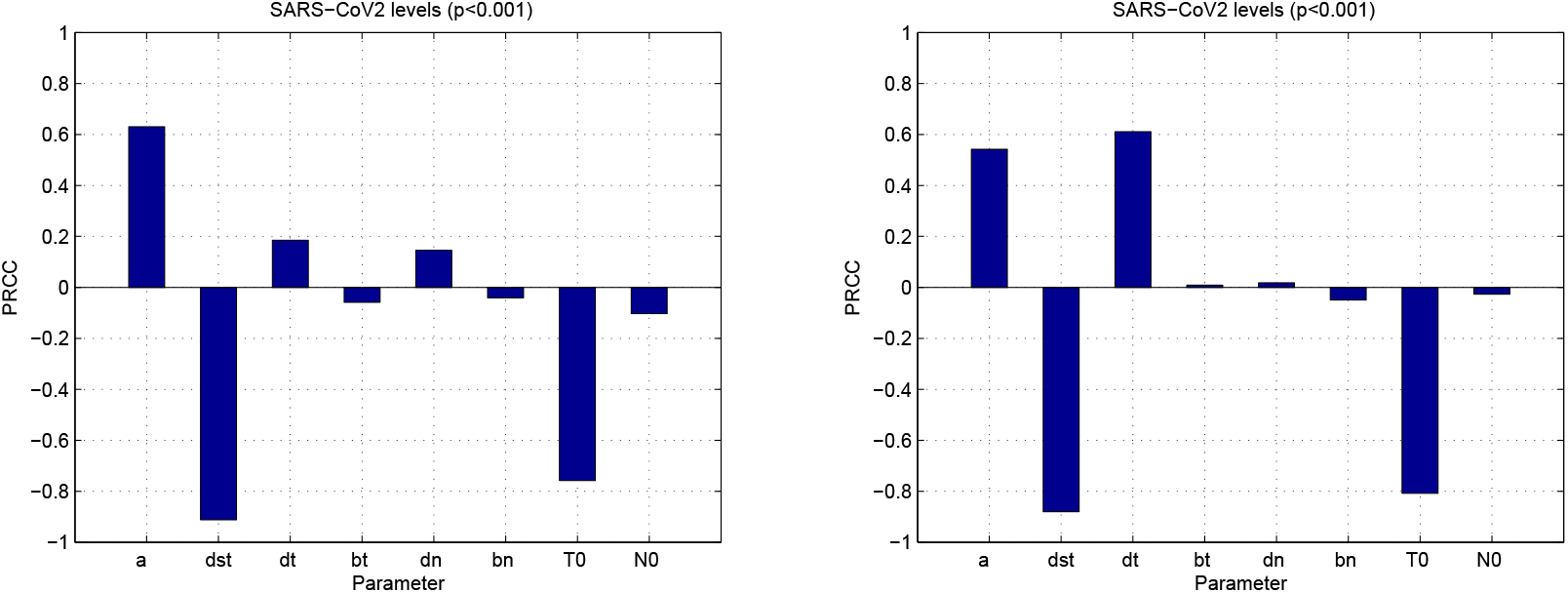
Sensitivity analysis for 4 (left) and 23 (right) days post-onset of symptoms.

On day 23 post-onset of symptoms, in addition to *d_st_* and *a*_1_ parameters, the *d_t_* parameter, which indicates the natural death rate of CD8^+^ T-cells, had a significant correlation with SARS-Cov-2. This correlation is positive with PRCC value 0.62, which indicates that in the forth week of the disease, death and consequently a decrease in the volume of cytotoxic T-cells has a great impact on the persistence of the virus and the disease is exacerbated.

Furthermore, to show the effect of *d_st_* and *d_t_* parameters on SARS-CoV-2 behavior in long COVID patients, we solved the model (1) with *α* = 0.8 for patient **g**, separately. According to Figure 11 (left), the maximum RNA viral for *d_st_* = 0.0275 is 1.2 × 10^7^ copies per ml, and the time for complete clearance of the virus is 330 days after the onset of symptoms. For *d_st_* = 0.0285 the maximum RNA viral is 5.2 × 10^6^ copies per ml, and the clearance time of the virus is 180 days after the onset of symptoms and for *d_st_* = 0.0295 maximum RNA viral and clearance time are 2.6 × 10^6^ copies per ml and 140 days after the onset of symptoms, respectively. As shown in Figure 11 (right) the maximum RNA viral for *d_t_* = 0.01 is 1.2 × 10^7^ copies per *ml*, and the time for complete clearance of the virus is 330 days after the onset of symptoms. So if we assume that vaccination increases the virus removal rate by CD8^+^ T-cells *d_st_* by 0.002, then vaccination of COVID-19 reduces the severity and effect of long COVID for 140 days. This is due to the induction of T cells with the vaccine.

**Figure 11:**
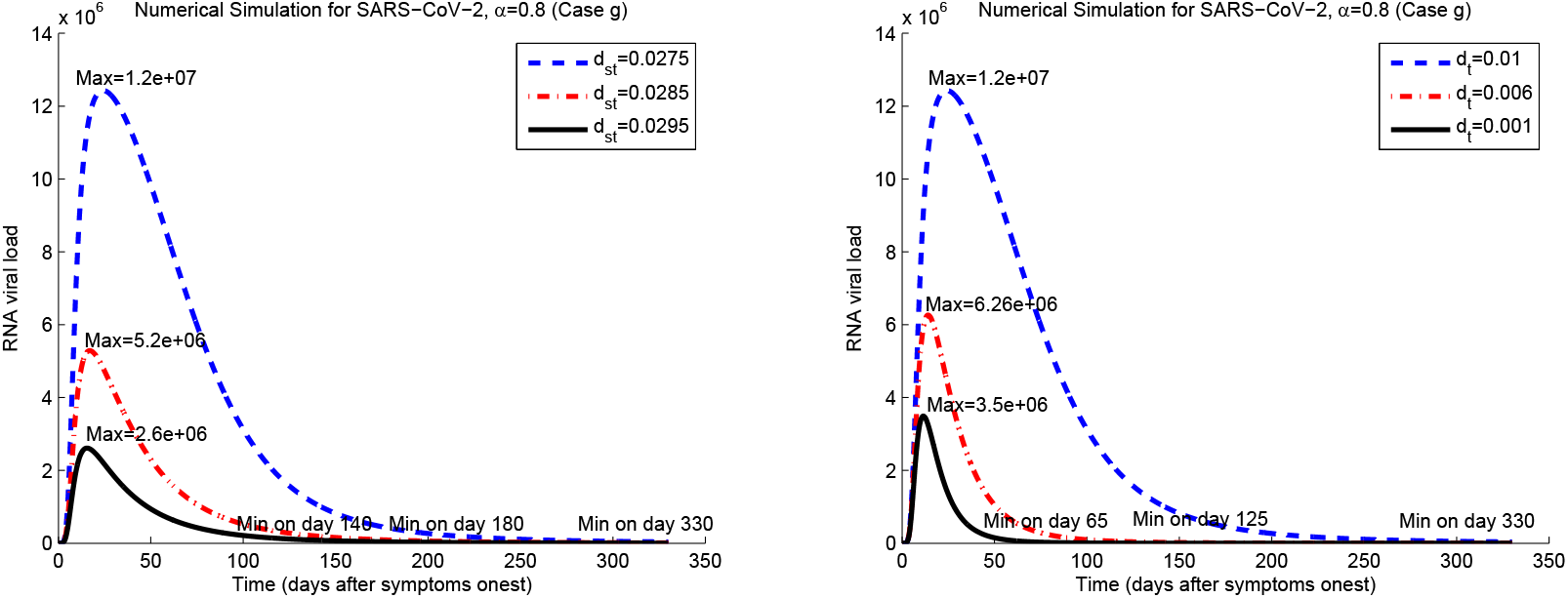
RNA viral load based on changing of *d_st_* (left) and *d_t_* (right) parameters for long COVID case **g**.

For *d_t_* = 0.006 the maximum RNA viral is 6.26 × 10^6^ copies per ml, and the clearance time of the virus is 120 days after the onset of symptoms and for *d_t_* = 0.001 maximum RNA viral and clearance time are 3.5 × 10^6^ copies per ml and 65 days after the onset of symptoms, respectively.

Thus, by increasing the lifespan of CD8^+^ T-cells by 0.005 and inducing long-term responses of these cells by vaccination, the long COVID period can be reduced to 65 days. Of note, this feature will be challenging for vaccine technology.

The findings published in [40] confirm the results of our model. In [40] it is shown that the symptoms and severity of long COVID among patients with persistent symptoms are significantly reduced 120 days after vaccination.

## 6 Conclusion

The coronavirus associated with severe acute respiratory syndrome-2 (SARS-CoV-2) interacts dynamically with many components of the immune system. These interactions are poorly understood because of their complexity. Using reliable mathematical models is one way to understand the mechanism of SARS-CoV-2 viral behavior. This paper presents a fractional-order mathematical model of the immune system responses to SARS-CoV-2 viral load in 5 patients with COVID-19. In this model, the population of cytotoxic T-cells (CD8^+^), natural killer cells are taken into account.

By sufficient conditions, non-negativity of the solution and asymptotic stability of the steady states are guaranteed. Simulation results shed light on the dynamics of SARS-Cov-2 and the immune system of the patients. Depending on the immune system, the dynamics of SARS-Cov-2 differ from person to person. It is possible for patients to develop so-called long COVID due to immuno-exhaustion. In Model 1, innate immunity, including NK cells, was well demonstrated. It is possible to achieve more results by developing the model and adding other parts of the immune system, such as helper T cells (CD4^+^).

A major advantage of the model was the fractional-order, which illustrated how age affects disease. In this case, the fractional-order value was 0 < *α* ≤ 1. Model (1) with *α* values closer to one is suitable for younger people and with smaller values is suitable for older people.

We performed a sensitivity analysis on some parameters to determine their effect on the model. SARS-Cov-2 load was closely correlated with some model parameters, such as the replication rate, virus removal rate by CD8+ T-cells, and death rate of T-cells. In addition to vaccine design, these parameters are useful in disease control and future treatments.

In the future, vaccine-related variables and parameters could be added to the model to prevent SARS-Cov-2 from spreading.

## Data Availability

Data are available on request from the corresponding author.

## References

[1] O. Diekmann, H. Heesterbeek and T. Britton, Mathematical tools for understanding infectious disease dynamics, Princeton University Press, 2012.

[2] WHO, Report of the WHO-China joint mission on coronavirus disease 2019 (COVID-19), World Health Organization, 2020.

[3] R. M. Anderson and R. May, Infectious diseases of humans: dynamics and control, Oxford, OUP, 1991.

[4] G. Marchuk, Mathematical modelling of immune response in infectious diseases, Dordrecht, Kluwer Academic Publishers, 1997.

[5] C. I. Siettos and L. Russo, Mathematical modeling of infectious disease dynamics,Virulence, 4(4) 295–306, (2013).

[6] W. Liu, H. W. Hethcote and S. A Levin, Dynamical behavior of epidemiological models with nonlinear incidence rates, J. Math. Biol., 25(4) 359–380, (1987).

[7] S. A Levin, B. Grenfell, A. Hastings and A. S. Perelson, Mathematical and computational challenges in population biology and ecosystems science, Science, 275(5298) 334–343, (1997).

[8] A. S. Perelson, Modelling viral and immune system dynamics, Natu. Rev. Immunol., 2(1) 28–36, (2002).

[9] A. S. Perelson and P. W. Nelson, Mathematical analysis of HIV-1 dynamics in vivo, SIAM Rev., 41(1) 3–4, (1999).

[10] E. M. Ahmed and H.A. El-Saka, On a fractional-order study of middle east respiratory syndrome coronalvirus (mers-cov), J. Fract. Calc. Appl., 8(1) 118–126, (2017).

[11] B. Yong and L. Owen, Dynamical transmission model of MERS-CoV in two areas, In AIP Conference Proceedings, volume 1716, pages 020010.1.020010.7, AIP Publishing, 2016.

[12] J. Lee, G. Chowell and E. Jung, A dynamic compartmental model for the middle east respiratory syndrome outbreak in the Republic of Korea: A retrospective analysis on control interventions and superspreading events, J. Theor. Biol., 408 118–126, (2016).

[13] A. A. Kilbas, H. M. Srivastava and J. J. Trujillo, Theory and applications of fractional differential equations in: North-Holland Mathematics Studies, 204, Elsevier Science B.V, Amsterdam, 2006.

[14] I. Podlubny, Fractional Differential Equations, Academic Press, USA, 1999.

[15] S. Ahmad, A. Ullah, Q. M. Al-Mdallal, H. Khan, K. Shah and A. Khan, fractional-order mathematical modeling of COVID-19 transmission, Chaos, Solitons and Fractals, 139 110256, (2020).

[16] K. Rajagopal, N. Hasanzadeh, F. Parastesh, I. I. Hamarash, S. Jafari and I. Hussain, A fractional-order model for the novel coronavirus (COVID-19) outbreak, Nonlinear Dyn., 101 711–718, (2020).

[17] H. Singh, H. M. Srivastava, Z. Hammouch and K. S. Nisar, Numerical simulation and stability analysis for the fractional-order dynamics of COVID-19, Results in Physics, 20 103722, (2021).

[18] I. A. Baba and B. A. Nasidi, fractional-order epidemic model for the dynamics of novel COVID-19, Alexandria Engineering Journal, 60 537–548, (2021).

[19] R. P. Yadav and R. Verma, A numerical simulation of fractional-order mathematical modeling of COVID-19 disease in case of Wuhan China, Chaos, Solitons and Fractals, 140 110124, (2020).

[20] A. N. Chatterjee, and B. Ahmad. A fractional-order differential equation model of COVID-19 infection of epithelial cells. Chaos, Solitons and Fractals, 147 (2021): 110952.

[21] A.N. Chatterjee, Fahad Al Basir, Muqrin A. Almuqrin, Jayanta Mondal, and Ilyas Khan. SARS-CoV-2 infection with lytic and non-lytic immune responses: A fractional order optimal control theoretical study. Results in physics 26 (2021): 104260.

[22] F. A. Rihan and G. Velmurugan, Dynamics and sensitivity analysis of fractional-order delay differential model for coronavirus infection, Progress in Fractional Differentiation and Applications, 7 43–61, (2021).

[23] J. Mondal, P. Samui, and A. N. Chatterjee. Optimal control strategies of non-pharmaceutical and pharmaceutical interventions for COVID-19 control. J. of Interdis. Math. (2020): 1–29.

[24] A. N. Chatterjee, F. Al Basir. A model for SARS-CoV-2 infection with treatment. Computational and mathematical methods in medicine. doi.org/10.1155/2020/1352982, (2020)

[25] F.A. Rihan, H.J. Alsakaji, Dynamics of a stochastic delay differential model for COVID-19 infection with asymptomatic infected and interacting peoples: Case study in the UAE, Results in Physics, 28(September), 2021, 104658.

[26] R. Wölfel et al., Virological assessment of hospitalized patients with COVID-2019. Nature, 581 465–469, (2020).

[27] S. Varchetta et al., Unique immunological profile in patients with COVID-19. Nature, Cell. and Mol. Immunol., 18 604–612, (2021).

[28] L. Mingyue, Elevated Exhaustion Levels of NK and CD8+T-cells as Indicators for Progression and Prognosis of COVID-19 Disease, Front. Immunol. doi: 10.3389/fimmu.2020.580237.eCollection, (2020).

[29] A. H. V. Esteban, In-host Mathematical Modelling of COVID-19 in Humans. Annu. Rev. in Con., 50 448–456, (2020).

[30] C. V. Eeden, Natural Killer Cell Dysfunction and Its Role in COVID-19, Int J. Mol. Sci., 21 6351, (2020).

[31] H. Mellstedt and A. Choudhur, T and B cells in in B-chronic lymphocytic leukaemia: Faust, mephistopheles and the pact with the devil, Cancer Immunology, 55, (2006) 210–220.

[32] A.Y Huang, P. Golumbek, M. Ahmadzadeh, E. Jaffee, D. Pardol, H. Levitsky, Role of bone marrow-derived cells in presenting MHC class I-restricted tumor antigens. Science 264(5161), (1994) 961–965.

[33] O. Diekmann, J.A. Heesterbeek, J.A., Metz, On the definition and the computation of the basic reproduction ratio R0 in models for infectious diseases. J. Math. Biol. 1990;35:503?522.

[34] M. Ye, J. Liu, C. Cenedese, Z. Sun and M. Cao, Network SIS meta-population model with transportation flow, IFAC PapersOnLine 53(2), (2020) 2562–2567.

[35] Z. M. Odibat and N. T. Shawagfeh, Generalized Taylors formula, Appl. Math. Comput., 186 286–293, (2007).

[36] L.G. de Pillis et al., A validated mathematical model of cell mediated immune response to tumor growth, Cancer Res., 65(1) 7950–7958, (2005).

[37] K. Diethelm et al., A predictor-corrector approach for the numerical solution of fractional differential equations, Nonlinear Dyn., 29 3–22, (2002).

[38] A. M. Baig, Chronic COVID syndrome: Need for an appropriate medical terminology for long-COVID and COVID long-haulers, J. of Medi. Virol., 93 2555–2556, (2021).

[39] S. M. Blower et al., Sensitivity and uncertainty analysis of complex models of disease transmission: An HIV model, as an example, Inter. Stati. Rev., 62 229–243, (1994).

[40] V-T. Tran, Efficacy of COVID-19 Vaccination on the Symptoms of Patients With Long COVID: A Target Trial Emulation Using Data From the ComPaRe e-Cohort in France. Preprints with The Lancet, https://papers.ssrn.com/sol3/papers.cfmabstract-id=3932953, (2021).

[41] L. Carolina, Kinetics of antibody responses dictate COVID-19 outcome, MedRxiv. Doi: 10.1101/2020.12.18.20248331. (2020).

[42] M. Vono et al., Robust innate responses to SARS-CoV-2 in children resolve faster than in adults without compromising adaptive immunity, Cell Rep. DOI: 10.1016/j.celrep.2021.109773, (2021).

[43] S. Nanda et al., B cell chronic lymphocytic leukemia: a model with immune response, Discrete Continuous Dyn. Syst. Ser. B, 18 1053–1076, (2013).

[44] S. Y. Zimmermann et al., A novel four-colour flow cytometric assay to determine natural killer cell or T-cell-mediated cellular cytotoxicity against leukemic cells in peripheral or bone marrow specimens containing greater than 20 percent of normal cells, J. Immuno. Meth., 296 63–76, (2005).

[45] R. J. DeBoer et al., Turnover rates of B cells, T-cells, and NK cells in simian immunodeciency virus-infected and uninfected rhesus macaques, The J. of Immuno., 170 2479–2487, (2003).

[46] A. M. Jamieson et al., Turnover and proliferation of NK cells in steady state and lymphopenic conditions, The J. of Immuno., 172 864–870, (2004).

[47] Y. Fadaei et al., A fractional-order mathematical model for Chronic Lymphocytic Leukemia and immune system interactions, Math Meth. Appl. Sci., DOI:10.1002/mma.6743, (2020).

[48] M. Hellerstein, M. Hanley, D. Cesar, S. Siler, C. Papageorgopoulos, E. Wieder, et al., Directly measured kinetics of circulating T lymphocytes in normal and HIV-1-infected humans, Nature Med., 5 83–89, (1999).

[49] A. Boukhouima, K. Hattaf and N. Yousfi, Dynamics of a fractional-order hiv infection model with specific functional response and cure rate, Int. J. Diff. Eqns, 2017 8372140, (2017).

[50] R. Garrappa. Predictor-corrector PECE method for fractional differential equations (https://www.mathworks.com/matlabcentral/fileexchange/32918-predictor-corrector-pece-method-for-fractional-differential-equations), MATLAB Central File Exchange. Retrieved December 17, 2021.

